# Zinc as a non-hormonal contraceptive: a better alternative to the copper intrauterine device (IUD)

**DOI:** 10.1101/2022.03.24.485705

**Authors:** Kirsten Shankie-Williams, Laura Lindsay, Chris Murphy, Samson Dowland

**Affiliations:** School of Medical Sciences, Faculty of Medicine and Health, The University of Sydney, Sydney, New South Wales, Australia

## Abstract

Long-acting and reversible contraceptives (LARC) are the most widely used form of female contraception worldwide, however they have significant side-effects that often result in early removal. Most LARCs are hormonal, but the use of exogenous hormones is not suitable for all women and causes side-effects in many others. The copper IUD (CuIUD) is the only non-hormonal LARC, but a large proportion of users suffer severe side effects. This study proposes the use of zinc as a suitable alternative to the CuIUD.

A rat intrauterine device (IUD) model was established to test the efficacy of a zinc IUD (ZnIUD) against a CuIUD, and a control nylon IUD. The IUD was surgically implanted into one uterine horn while the other remained untreated. Both the ZnIUD and CuIUD resulted in significantly fewer implantation sites compared to untreated horns. There was no significant difference between treated and untreated horns in the control nylon IUD group. Histological assessment revealed damage and inflammation in the endometrium of CuIUD treated horns, but only minor epithelial damage in ZnIUD treated horns, closely resembling the normal appearance of the control horns. This suggests ZnIUDs may not share the side-effect profile of the CuIUD. To test the long-term efficacy of the ZnIUD, rats had a ZnIUD surgically implanted into both horns and cohoused with males for 3 months. These rats mated regularly but did not get pregnant, confirming the long-term effectiveness of the ZnIUD. Reversibility of the ZnIUD was also established, as removal of the ZnIUD after 3 months resulted in no significant difference in the number of implantation sites between treated and untreated horns.

This study demonstrated the contraceptive efficacy of zinc and its potential as a LARC. The ZnIUD had minimal histological impact on the endometrium compared to the current copper standard, indicating that IUDs containing zinc may offer highly effective contraception while causing fewer side effects.

## Introduction

Intrauterine devices (IUDs) are the most commonly used form of reversible contraception world-wide (1). Almost all long-acting reversible contraceptives (LARCs) contain hormones as their contraceptive agent and many women are either precluded from using these by a medical condition or wish to avoid the many physical and physiological side-effects of exogenous hormones. Depression, mood swings, irritability, anxiety, weight gain, migraines and decreased libido are just some of the commonly reported side-effects experienced by hormonal contraceptive users in numerous studies over the last few decades (2–6). These can lead to discontinuation or inconsistent use, both increasing the chance of unplanned pregnancies. Only one non-hormonal option currently exists, the copper IUD (CuIUD), which is a medicated model that when inserted into the uterus provides the user up to 12 years of protection against pregnancy. This long-term contraceptive comes with many practical benefits distinguishing it from alternatives, such as the contraceptive pill, including low cost, no potential for patient error and no exogenous hormone induced side-effects. These characteristics provide the foundation for a low risk and financially accessible contraceptive option for women around the world. However, the CuIUD has its own unique side-effect profile causing more painful periods for 38% of women (7). This excessive pain and bleeding, both menstrual and intermenstrual, has led to persistent rates of early removal of around 25%. For this reason it is not recommended for women with moderate to heavy periods (7). 1 in 10 women worldwide have an unmet need for contraception and side-effects are a primary reason women are left with no protection, resulting in 121 million unintended pregnancies each year (8–10).

Side-effects experienced by users of the CuIUD are a direct result of the copper content of the device. Animal models have shown that copper cytotoxicity causes necrosis, erosion, stromal hypertrophy and metaplasia of epithelial cells in mucosal surfaces in direct contact with copper, resulting in polyps (11,12). Exposure to metallic copper also resulted in an increase in lysosomes and swollen, vesiculated mitochondria in vaginal epithelial cells and infiltration of leukocytes into the underlying mucosa (12). This is consistent with the morphological changes observed in human endometrial biopsies taken from women using the CuIUD (13). Lymphocytic, monocytic and leukocytic infiltrations have been found in the uterine epithelium and stroma of copper IUD users, and a loss of epithelial integrity was observed (14).

During early experiments exploring suitable non-hormonal IUD materials a range of metals were trialled for antifertility effects. The first study to discover the antifertility effects of copper tested the contraceptive actions of copper, zinc, silver, tin and magnesium in the rabbit uterus by placing an IUD of each metal near the cervix (11). Copper and zinc resulted in the highest anti-fertility rates of all the metals studied, finding 1 implantation site across 6 animals treated with zinc and no implantation sites in the copper treated animals (11). After early studies showed a slightly lower contraceptive efficacy of zinc than copper, further investigation of the use of zinc in an IUD was largely abandoned. However, we suggest that the slightly lower efficacy of zinc in this study may have resulted from methodological limitations. Although the use of zinc as a contraceptive was ultimately not pursued much further, later studies have shown that the 2-cell embryo is uniquely vulnerable to zinc and exposure of the 2-cell embryo to zinc significantly reduces blastocyst formation (15).We therefore hypothesise that zinc can act on the 2-cell embryo to exert a contraceptive effect and that if this mechanism is taken into consideration, a ZnIUD could be an effective LARC. This indicates that copper and zinc produce a contraceptive effect via different mechanisms of action. As such, the placement of the experimental IUDs near the cervix of rabbits in the Zipper et al. (11) study was suitable for copper, but suboptimal for zinc – being located distant from the site at which we now expect it to act.

A 1972 study by Medel et al. (16) claimed to demonstrate the reversibility of copper and zinc IUDs in rats and rabbits. This is the only previous study to examine zinc IUD reversibility, but it contains methodological limitations/issues that limit the validity of the findings as claimed. This model involved animals being mated 2-3 days after insertion of IUDs, which were then removed on day 2 of pregnancy, with pregnancy outcomes evaluated on day 10 (16). This work showed pregnancy was able to proceed in these animals; however, this study design only evaluates the impact of IUD removal in early pregnancy rather than demonstrating reversibility of these contraceptives as claimed. The study contains multiple potentially confounding factors, in particular performing surgery on the uterus during pregnancy could impact the success and final number of implantation sites. Physiological responses to anaesthesia including decreased heart rate and body temperature could affect implantation success or maintenance of implanted blastocysts (17). Moreover, as the authors acknowledge, surgical trauma to the uterus during pregnancy would most certainly contribute to the reported findings of decreased fertility in the IUD treated horns.

Although these earlier studies provided initial evidence supportive of zinc as a contraceptive agent; true efficacy and reversibility of a ZnIUD have not yet been established. Therefore, the present study has developed a rat model to examine the contraceptive efficacy of a zinc IUD with consideration of its potential mechanism of action. This paper also examines the potential for zinc to provide continued contraception over an extended period of time and assesses the return to fertility after such long-term use of a ZnIUD, to determine the true reversibility of such a contraceptive. The impact of a ZnIUD on the endometrium is also compared with that of a CuIUD. This work aims to evaluate the potential utility of a ZnIUD as a non-hormonal LARC, in comparison with the only current option, the CuIUD.

## Methods

### Animals and mating

23 nulliparous female Wistar rats were housed in plastic cages under a 12 hr light-dark cycle at 21°C. All rats were between 10-12 weeks old at the start of the protocol and weighed 280-300g. Rats were fed and watered *ad libitum*. All procedures were approved by The University of Sydney Animal Ethics Committee. The oestrous cycle of the rats was tracked by vaginal smearing and analysis of the cellular composition (18). Mating was also confirmed by smearing, the presence of sperm indicating day 1 of pregnancy. Pseudopregnancy was confirmed by the cellular composition of daily vaginal smears (18).

### Treatment groups

Rats were randomly allocated to 3 experimental groups (Fig. 1). In the Efficacy Group, rats had either a zinc (n= 5), copper (n= 6) or nylon IUD (n= 5) inserted into one uterine horn, were mated after recovery and tissue was collected on day 9 of pregnancy. In the Long-term Group (n=3), rats had zinc IUDs inserted into both uterine horns, allowed to recover for 1 week then co-housed with a male for 3 months. Rats were regularly smeared to track the oestrous cycle, mating events, and number of days in pseudopregnancy. At the end of the 3-month period tissue was collected 9 days after mating. In the Reversibility Group (n= 4), rats had a zinc IUD inserted into one uterine horn which was removed 3 months later. After recovery, animals were mated, and tissue was collected on day 9 of pregnancy.

**Figure 1.**
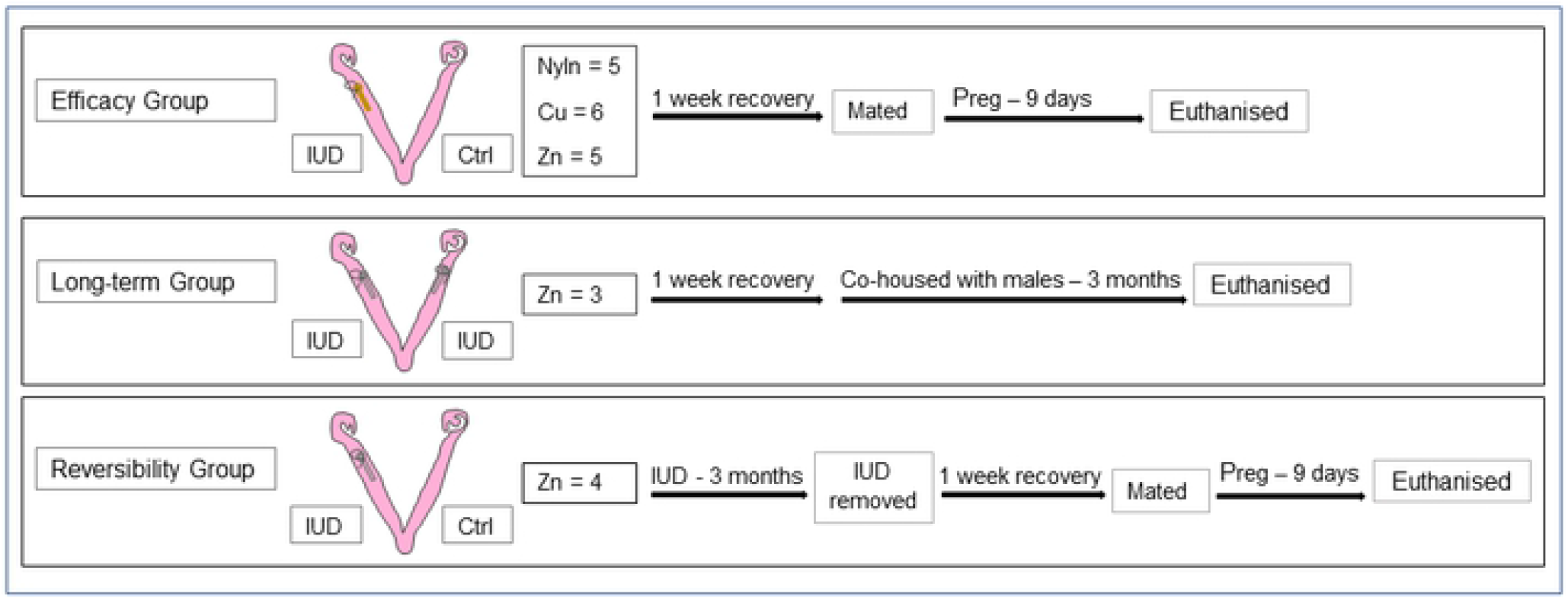
Flowchart of experimental treatment groups. The Efficacy Group had either a zinc (n= 5), copper (n= 6) or nylon IUD (n= 5) inserted into one uterine horn. The Long-term Group (n=3) rats had zinc IUDs inserted into both uterine horns for 3 months. The Reversibility Group (n= 4) had a zinc IUD inserted into one uterine horn which was removed 3 months later. Animals were mated and tissue was collected on day 9 of pregnancy.

### Surgical Insertion and Removal of IUDs

#### Insertion

Zinc and copper IUDs were made from 1 cm of 99.994% pure zinc and 99.999% pure copper wire (Alfa Aesar, MA, USA), 0.5 mm in diameter, and were finished with a small loop at one end. Rats received an intraperitoneal injection of ketamine (75mg/kg; TROY Laboratories, NSW, Australia) and xylazine (10mg/kg; Parrell Laboratories, NSW, Australia). IUDs were inserted into the uterine lumen through a small incision in the uterine wall, 1 cm from the ovary. All IUDs were secured in place with a nylon suture through the IUD loop and uterine wall. A foreign body control was also performed by inserting a 1cm loop of nylon thread into the uterine horn to test the effect of the presence of a foreign object in the uterine lumen. Rats were allowed to recover for at least one week after surgery.

#### Removal

Removal surgery for animals in the Reversibility group occurred during dioestrus, at which time the uterus is small, not oedematous and lacks prominent vasculature, allowing easier removal and minimising tissue damage. Similar to insertion surgery, rats received an intraperitoneal injection of ketamine (75mg/kg) and xylazine (10mg/kg), the uterine horn was exposed, and the IUD was carefully removed. Rats recovered for at least a week.

### Tissue collection

Rats were anaesthetised with 20mg/kg sodium pentobarbitone (TROY Laboratories, NSW, Australia) and dissected under deep anaesthesia before euthanasia. Both uterine horns were excised, pinned onto dental wax and photographed. The implantation sites were counted and compared to the control horn to determine contraceptive efficacy (19). Uteri were cut into approx. 5mm blocks from the region of the horn containing the IUD or inter-implantation sites in the control (20) and immediately fixed in 10% neutral buffered formalin (NBF; Fronine, NSW, Australia). Tissue was then washed in distilled H_2_O and dehydrated in 30, 50 and 70% ethanol for 30 minutes each. Samples were then placed in the Excelsior ES tissue processor (Thermo Scientific, USA) to further dehydrate tissue in graded ethanol baths over 4.5 hrs at 40°C before clearing with two 45-minute changes of xylene and infiltration with paraffin (Thermo Fisher, USA) over 3 changes of 30 minutes at 60°C. Finally, specimens were embedded using the Tissue-Tek TEC 5 tissue embedding console system (Sakura FineTek, USA).

### Sectioning and Staining

Transverse sections (7μm) were cut on an HM 325 Rotary Microtome (Thermofisher Scientific), sections floated on a water bath (40°C) and placed onto gelatin chrom-alum coated glass slides. At least 5 sections were collected per block and 2 blocks were used for each animal. Slides were then dried overnight in a 40°C oven.

Sections were stained using a routine haematoxylin and eosin protocol (21). Images were captured on an Olympus BX53 brightfield microscope (Olympus, VIC, Australia), using an Olympus DP73 camera (Olympus) and Olympus CellSens software (version 1.11).

### Statistical analysis

In the Efficacy Group changes in the number of implantation sites in IUD treated horns and contralateral non-treated horns were analysed using two-way analysis of variance (ANOVA), with IUD material and uterine horn as factors. Tukey post hoc test for multiple comparisons (reporting multiplicity-adjusted P values) was then applied to determine which pairs of means were significantly different. Differences were determined to be statistically significant when P < .05. For the data obtained in the Reversibility Group a paired *t*-test was used to determine if a statistical difference existed between the number of implantation sites present in a horn that had a ZnIUD removed after 3 months and a contralateral non-treated horn. A *p*-value of ≤ 0.05 was considered statistically significant. All statistical analysis was performed using GraphPad Prism Software (version 7.02; GraphPad Software, Inc, La Jolla, California) and bar graphs were generated where values represent the mean + SEM.

## Results

### Copper and Zinc IUDs prevent pregnancy

The contraceptive efficacy of copper and zinc was assessed by counting the number of implantation sites in the IUD treated and untreated horns on day 9 of pregnancy in rats receiving different IUD materials (Fig. 2). In rats receiving a nylon IUD, implantation sites were evident in both the IUD horn and untreated, control horn with no significant difference in the number of implantation sites in control vs treated horns (P= 0.1059) (Fig. 2). In rats with either a CuIUD or ZnIUD there were no implantation sites in uterine horns containing an IUD. Implantation sites were evident in the untreated uterine horns, resulting in a statistically significant difference in the number of implantation sites in treated compared to untreated horns in both CuIUD (P<0.0001) and ZnIUD (P<0.0001) animals (Fig. 2).

**Figure 2.**
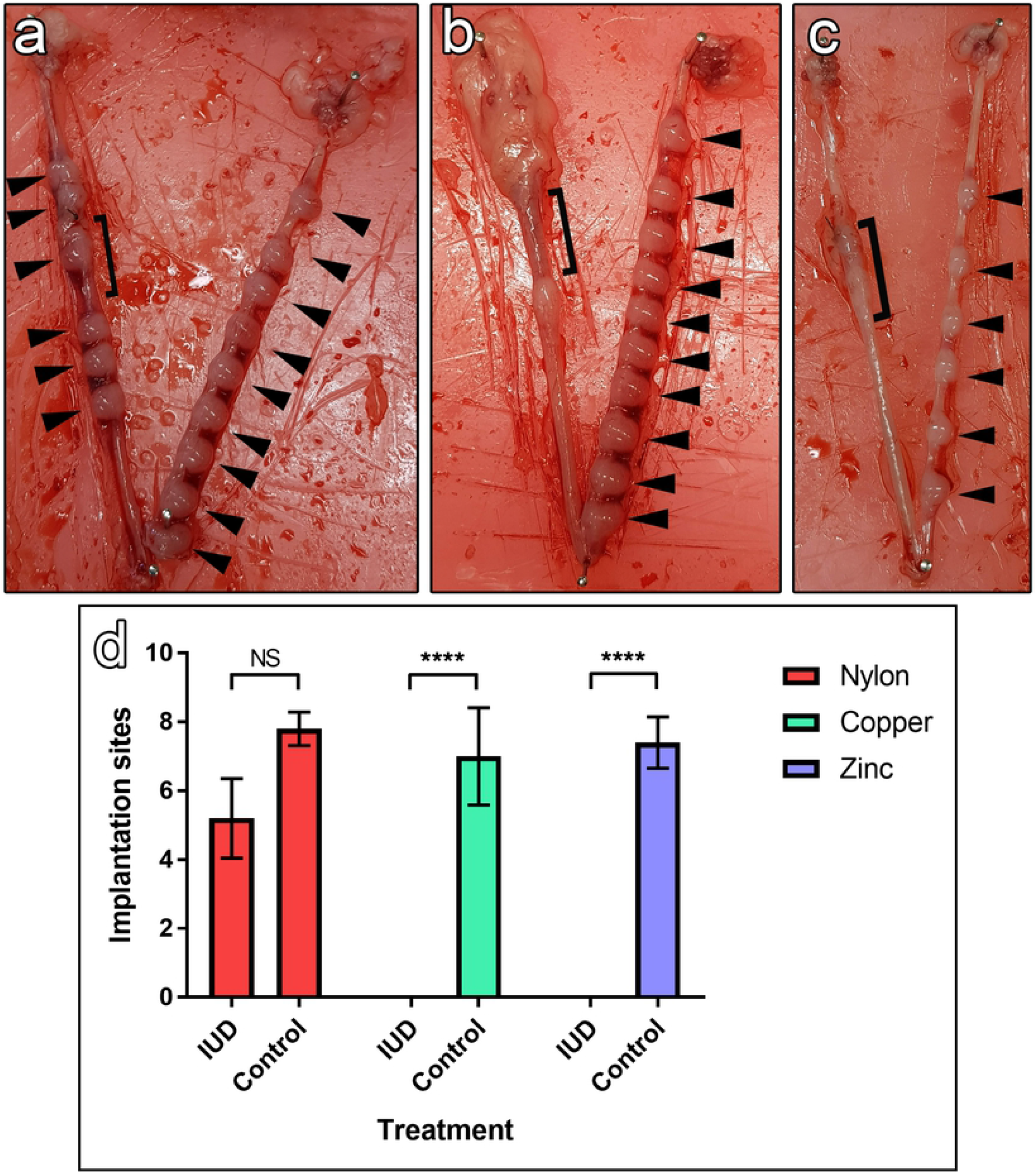
Contraceptive efficacy of metal IUDs (Cu^2+^ + Zn^2+^) and a control nylon IUD on day 9 of pregnancy. **(a)** Implantation sites (arrow heads) were visible in both IUD and control horns of the nylon treated rats, as well as the region containing the IUD (bracket). **(b)** CuIUD treated horns did not contain implantation sites while the non-treated control horn contained multiple. **(c)** ZnIUD treated horns did not contain implantation sites while the non-treated control horn contained multiple. **(d)** There was no significant difference in the number of implantation sites between treated and control uterine horns in rats receiving a nylon IUD (n= 5). No implantation sites were seen in the CuIUD (n= 6) and ZnIUD (n=5) treated uterine horns in any animals, while the respective control horns contained implantation sites. (**** = P<0.0001).

### Short-term Exposure to Metal IUDs Impacts Endometrial Histology

Endometrial histology was assessed after exposure to each of the IUD materials. Uterine horns containing a nylon IUD exhibited no abnormalities and displayed normal uterine morphology (Fig. 3a). The glands and myometrium appeared normal. The untreated horn demonstrated normal histological appearance with no signs of damage, inflammation or metaplasia. The endometrium of the copper treated uterine horn contained sites of epithelial damage in the region of the IUD (Fig. 3b). The adjacent uterine stroma exhibited infiltration of inflammatory cells, including dense clusters of macrophages and lymphocytes (22). Along areas of intact epithelium, the simple columnar epithelial cells had undergone metaplasia with a resulting saw-tooth appearance. The underlying stroma, glands and myometrium of the uterus appeared normal. The untreated uterine horn demonstrated normal histological appearance of the endometrium with no epithelial damage, inflammation, or metaplasia. In the zinc IUD-treated uterine horns the epithelium showed mild metaplasia resulting in an uneven apical border (Fig. 3c). The epithelium remained intact, with no regions of epithelial damage or exposure of the stroma and no inflammation was present. The underlying stroma, glands and myometrium appeared normal. The untreated uterine horn did not show signs of damage, inflammation, or metaplasia (Fig. 3d).

**Figure 3.**
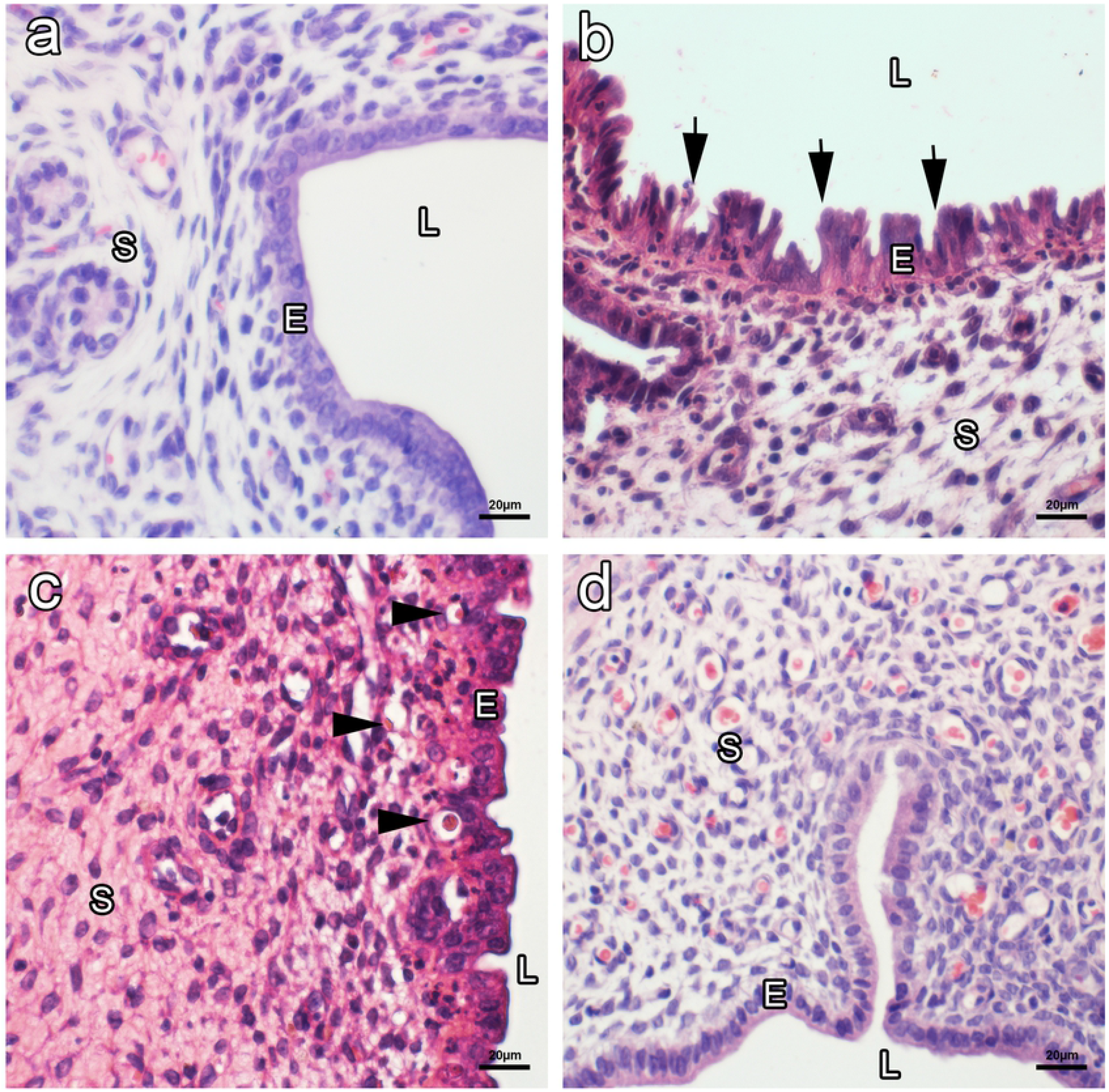
Uterine histology of horns containing metal IUDs (Cu^2+^ and Zn^2+^), a control nylon IUD and the contralateral non-treated horn on day 9 of pregnancy. **(a)** Haematoxylin and eosin showing no abnormalities present in the nylon IUD treated horn. **(b)** Metaplasia (arrows) of the uterine epithelial cells is present in the CuIUD treated horn. **(c)** Mild metaplasia of the uterine epithelial cells is present in the ZnIUD treated horn and many capillaries are apparent (arrow head). **(d)** The contralateral non-treated horn exhibits normal uterine morphology.

### The Zinc IUD Provides Long-term Contraception

The long-term contraceptive efficacy of a ZnIUD was assessed by inserting IUDs in both uterine horns and co-housing rats with males for 3 months. Daily vaginal smears showed that the rats regularly mated but did not get pregnant. Cervical stimulation during mating causes rescue of the corpus luteum in rats resulting in rats spending an average of 10 days in a pseudopregnant state before returning to oestrus and mating again (23). Despite rats mating at least 8 times each, there were no pregnancies during the 3-month treatment period (Table 1).

**Table 1.**
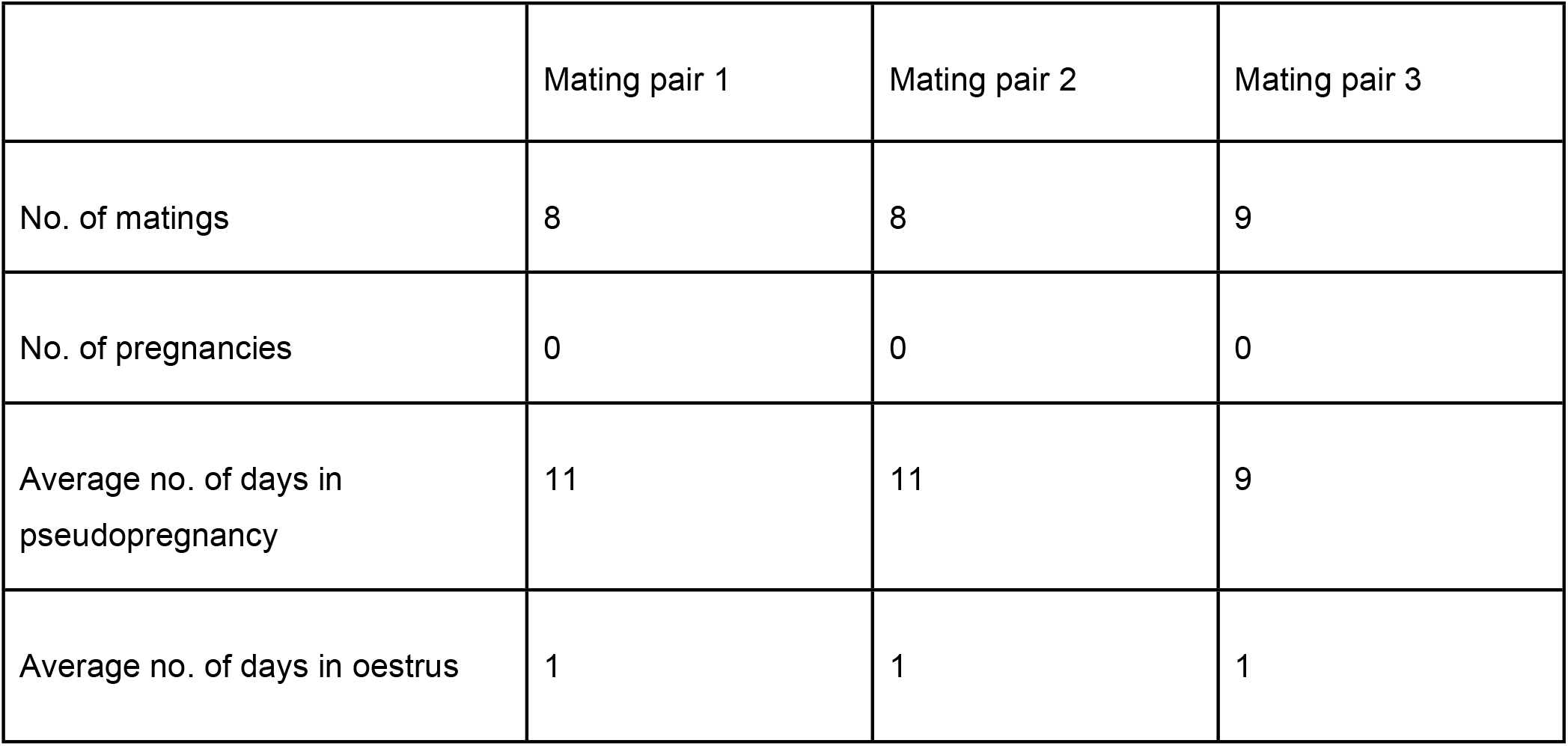
Contraceptive efficacy of the ZnIUD over a 3 month period

|  | Mating pair 1 | Mating pair 2 | Mating pair 3 |
| --- | --- | --- | --- |
| No. of matings | 8 | 8 | 9 |
| No. of pregnancies | 0 | 0 | 0 |
| Average no. of days in pseudopregnancy | 11 | 11 | 9 |
| Average no. of days in oestrus | 1 | 1 | 1 |

### The Contraceptive Effect of the Zinc IUD is Reversible

The reversibility of the method was investigated by inserting a ZnIUD into one uterine horn, removing it 3 months later then examining implantation success. No significant difference in the number of implantation sites in the ZnIUD treated uterine horn compared to the untreated horn was found (P= 0.86)(Fig. 4). The average number of implantation sites in ZnIUD treated uterine horns was 5.75 (±1.4) compared to 6 (±0.4) implantation sites in control horns.

**Figure 4.**
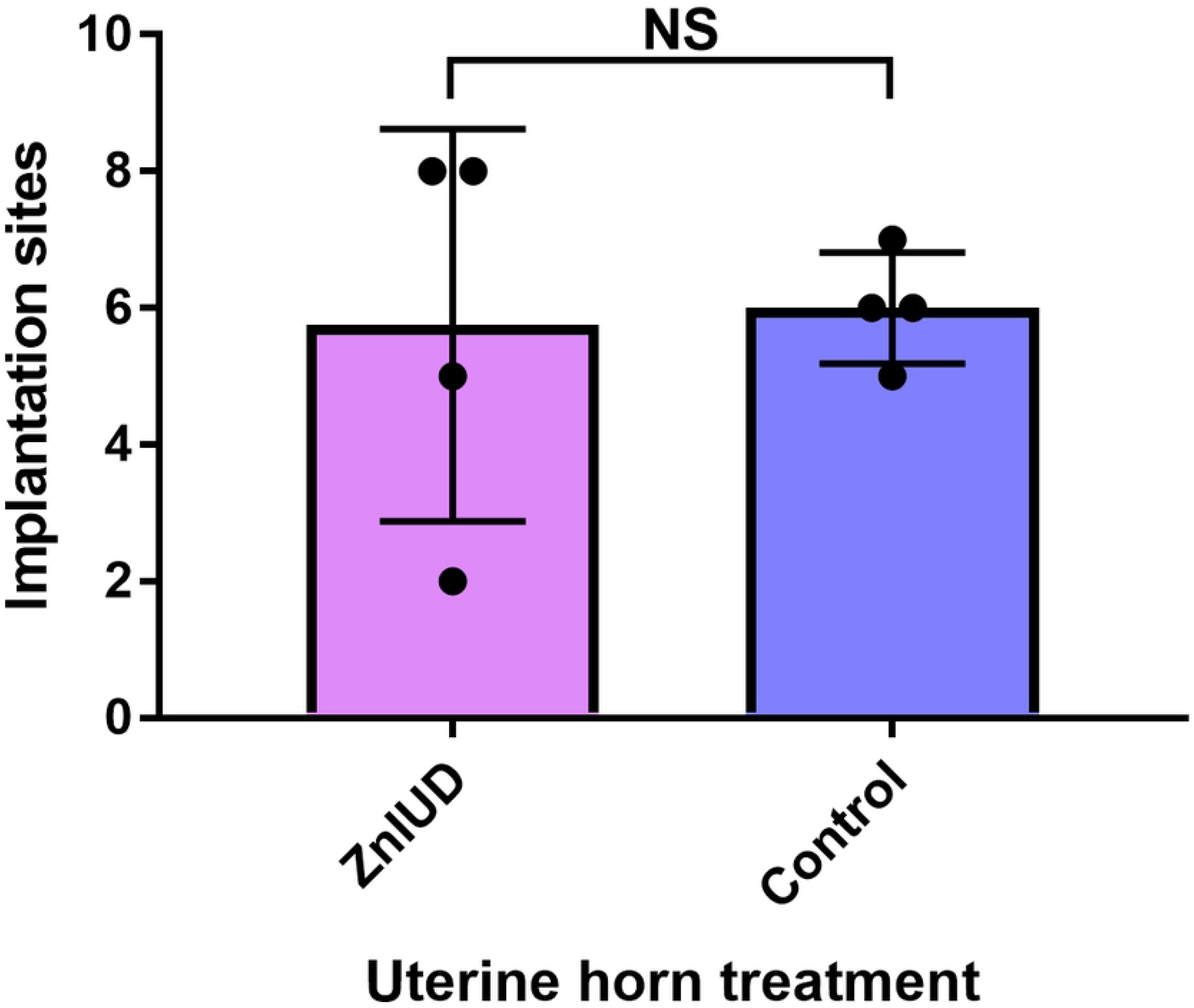
Number of implantation sites in uterine horns after a 3-month treatment and subsequent removal of a zinc IUD (n=4). There was no significant change in implantation rate between treated and untreated horns after IUD removal from the treated horns.

**Figure 5.**
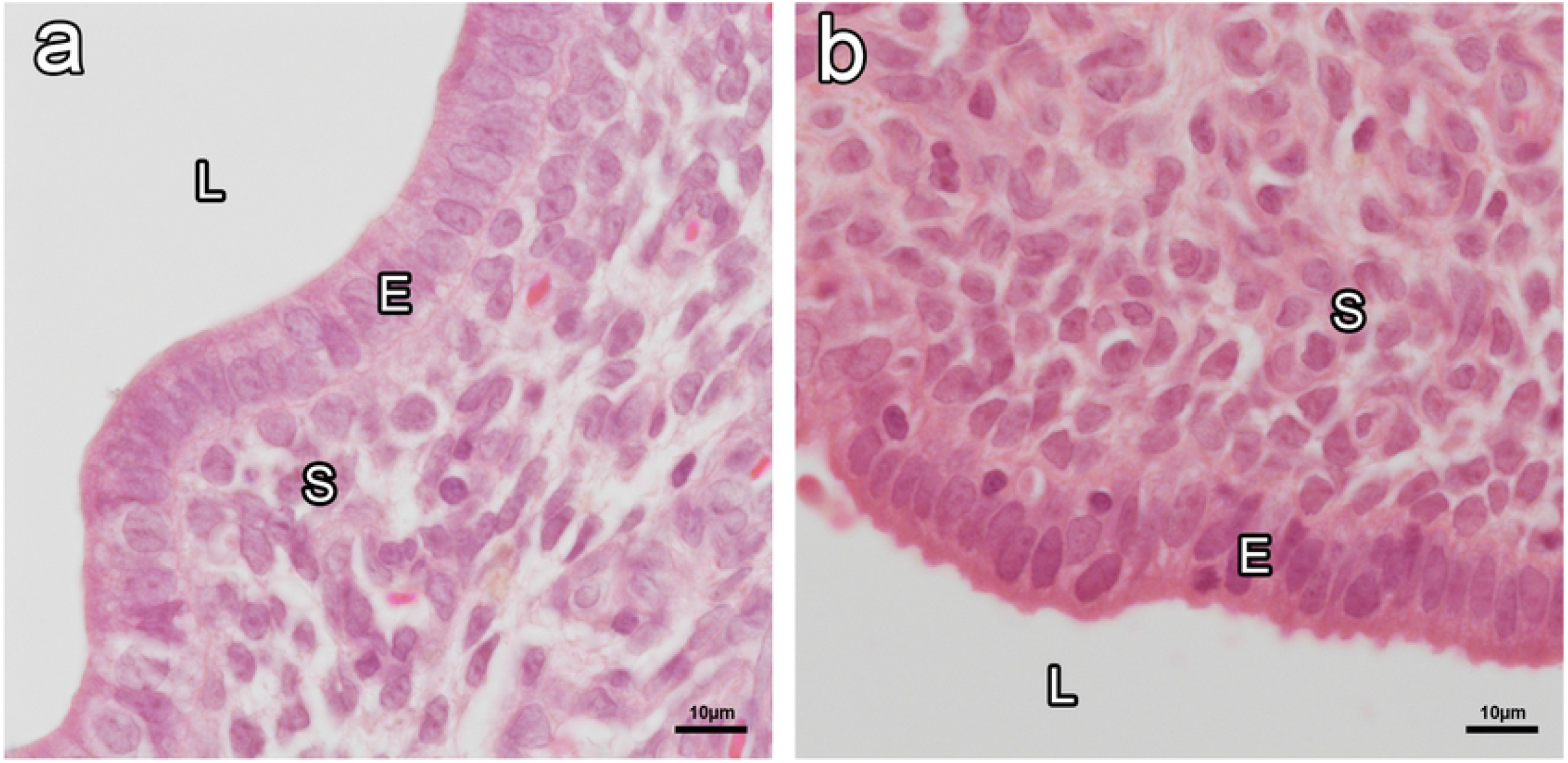

### Long-term Exposure to the Zinc IUD does not Impact Endometrial Histology

Histological analysis of the endometrium after 3 months exposure to a ZnIUD revealed no pathologies such as metaplasia or inflammation in the ZnIUD treated uterus. The untreated horn did not show signs of damage, inflammation or metaplasia.

## Discussion

Non-hormonal long-acting reversible contraceptives are cheap and effective however the current CuIUD has significant side-effects that limit its uptake. This study used a rat IUD model to investigate an alternative ZuIUD for its contraceptive efficacy, reversibility and long-term reliability.

### Both Copper and Zinc demonstrate 100% Contraceptive Efficacy in the Rat IUD Model

In agreement with previous animal studies, the present study determined that the CuIUD provided 100% contraceptive efficacy, completely preventing pregnancy in the IUD-treated uterine horns. This study also demonstrated for the first time that a ZnIUD can provide 100% contraceptive efficacy, equal to that of the CuIUD. Comparison with the nylon IUD demonstrated that the contraceptive effect of both the ZnIUD and CuIUD is due to the metal content of the IUD, rather than resulting from the presence of a foreign body in the uterine lumen. Our IUD model demonstrated higher contraceptive efficacy than Zipper et al. (11), showing this model successfully takes into consideration the mechanism of action.

### The Zinc IUD is an Effective Long-term and Reversible Contraceptive in the Rat

In order to mimic the clinical setting, this study for the first time showed the long-term contraceptive efficacy and reversibility of a ZnIUD. The ZnIUD prevented all pregnancies in female rats co-housed with males for a continuous 3 month period with no change in the number of days in oestrus or the average days of pseudopregnancy after cervical stimulation (18). This indicates that zinc may have long-term efficacy in clinical settings similar to that of the CuIUD. The present study is also the first to clearly demonstrate that the contraceptive effect of a ZnIUD is reversible. When ZnIUDs were removed after 3 months exposure, rats were able to rapidly return to fertility and exhibited successful implantation in uterine horns previously containing an IUD, at rates comparable to untreated uterine horns. This is in contrast to the previous study which found a significant decrease in implantation sites when zinc IUDs were removed on the second day of pregnancy, in treated compared to control horns (16). As such, this study has shown that the ZnIUD could be an effective LARC and offer an alternative contraceptive option to women.

### The Copper IUD Impacts Endometrial Histology More Severely than Zinc

The morphological changes and pathologies of the CuIUD treated horn, observed histologically in the present study, reflect those seen in the earlier rabbit and human trial of the copper IUD (11). The CuIUD in our rat IUD model caused severe metaplasia in the uterine epithelium and epithelial damage. Inflammation was also present in the CuIUD horn, in alignment with the stromal hypertrophy documented by Zipper et al. (11) in copper treated rabbit uterus (11). An influx of inflammatory cells around areas that had lost epithelial integrity were common amongst the copper treated horns. This inflammation and endometrial damage may lead to many commonly reported side-effects. Inflammatory mediators activate and sensitise local nociceptors to prevent further tissue damage (24), whilst the continuous release of proteolytic agents from leukocytes suppress endometrial angiogenesis, resulting in pain and excess bleeding (25). Recent studies have found a higher concentration of copper in the endometrium of women experiencing abnormal bleeding with a CuIUD and a positive correlation with VEGF, a potent angiogenic factor (26). In addition, higher fibrinolytic activity has been found in areas of the endometrium that have direct contact to the CuIUD, indicating inhibition of homeostasis (27). The histological data in the present study supports these observations, further supporting that copper directly exerts negative effects on the endometrium.

These effects are particularly severe soon after insertion, when the rate of dissolution of copper ions into the uterine fluid is highest upon insertion of the CuIUD and drops exponentially over the duration of its use, an effect known as the ‘burst release’ (28). This high initial copper corrosion rate irritates the endometrium, enhancing the inflammatory response and deposition of proteins and cell debris and may contribute to the inflammation and damage of the endometrium after short-term exposure to the CuIUD, seen in this study. This period is when the highest removal rates occur, preventing many women from having successful, long-term use of the CuIUD (7).

The impact of a ZnIUD on uterine histology has not been previously examined. Short-term exposure (approx. 2-3 weeks; Efficacy Group) to the ZnIUD resulted in mild metaplasia in the uterine horn, with epithelial cells appearing irregular in height. Unlike the CuIUD treated horn the epithelium remained intact, and no areas of inflammation were observed. Therefore, it is possible that a ZnIUD would cause users less pain and bleeding compared to the CuIUD, due to the absence of localised inflammation (24,25). Uterine horns that were subjected to long-term exposure to Zn (approx. 16 weeks; Long-Term Group) exhibited no pathologies, having the same appearance as the control horn. This indicates that zinc acts as a contraceptive without any long-term impact to endometrial integrity. Although there have been significantly fewer studies conducted on the interaction of zinc in the uterus compared to those of copper, the present findings, in conjunction with the emerging use of zinc in biocompatible implants provides a positive indication of the potential biocompatibility of a ZnIUD (29,30). Zinc is relatively harmless to humans as it’s the second most abundant essential metal in the body and found at a mean serum concentration of 1000μg/ml in humans, with a recommended daily intake of 10-15mg (31). This is in contrast to copper, which is found at a mean serum concentration of 0.1μg/ml, and a recommended daily intake of 0.9mg. Because of this, the body is physiologically adept at transporting and storing larger concentrations of zinc with no risk of producing reactive oxygen species, a consequence of high concentrations of copper and mediators of acute and chronic inflammatory pain (32,33). Zinc is now recognised as an excellent candidate for zinc-based orthopaedic implants, being chosen for its biocompatibility and safety (30,34).

### The contraceptive effect of a Zinc IUD is likely via a Unique Mechanism, Acting on the 2-Cell Embryo

The improved contraceptive efficacy of zinc in the present study compared to earlier studies (11,35,36), supports our hypothesis that zinc exerts its contraceptive effect via a unique effect on the 2-cell embryo. Previous research has shown that the 2-cell embryo exhibits unique sensitivity to Zn^2+^ (15). At this stage the embryo has begun transcription of metallothionein, an important metal binding protein. However, at such an early stage metallothionein is a precursor protein without metal reactivity (15). The metal responsiveness occurs later in development meaning the presence of the slightly increased concentrations of Zn2+ during this window of development may be lethal to the embryo. The location of the ZnIUD prior to fertilisation in our model facilitates high contraceptive efficacy and would explain the lack of a contraceptive effect seen in earlier studies when a zinc wire was inserted 3 days post fertilisation (35), when embryos would already have progressed past the 2-cell stage. The present study also demonstrates higher contraceptive efficacy than another model used by Kincl and Rudel (36), where a 3mm zinc loop was inserted into the uterus 3 days post copulation. This supports our suggestion that zinc acts on the 2-cell embryo, which is uniquely vulnerable to zinc (15), such that the ZnIUD must be in place within 24hrs post-fertilisation, before the zygote cleaves into 2 cells.

This study has refined an animal model to study the effectiveness of non-hormonal IUDs in the rodent uterus and for the first time allowed direct evaluation of the endometrial responses to these metals. This model showed that a ZnIUD provides long-term and reversible contraception without causing local tissue damage to the endometrium. As such the ZnIUD stands up to the gold-standard for non-hormonal contraception, the CuIUD. The ZnIUD may also have an additional advantage of a new mechanism of action that impacts the embryo directly, avoiding irritation and inflammation of the uterus. Together, these findings suggest that the ZnIUD may have utility as a LARC that could have significantly fewer side-effects.

## Acknowledgements

This work was supported by an NWG Macintosh Memorial Grant from The Ann Macintosh Foundation of the Discipline of Anatomy and Histology and Murphy Laboratory Funds.

